# Algorithm for automatic detection of insulin granule exocytosis in human pancreatic β-cells

**DOI:** 10.1101/2023.11.14.566999

**Authors:** Aishwarya A Makam, Abhimanyu Dubey, Shovamayee Maharana, Nikhil R. Gandasi

## Abstract

Image processing and analysis are two significant areas that are highly important for interpreting enormous amounts of data obtained from microscopy-based experiments. Several image analysis tools exist for the general detection of fundamental cellular processes, but tools to detect highly distinct cellular functions are few. One such process is exocytosis, which involves the release of vesicular content out of the cell. The size of the vesicles and the inherent differences in the imaging parameters demand specific analysis platforms for detecting exocytosis. In this direction, we have developed an image-processing algorithm based on Lagrangian particle tracking. The tool was developed to ensure that there is efficient detection of punctate structures initially developed by mathematical equations, fluorescent beads and cellular images with fluorescently labelled vesicles that can exocytose. The detection of these punctate structures using the tool was compared with other existing tools, such as find maxima in ImageJ and manual detection. The tool not only met the precision of existing solutions but also expedited the process, resulting in a more time-efficient solution. During exocytosis, there is a sudden increase in the intensity of the fluorescently labelled vesicles that look like punctate structures. The algorithm precisely locates the vesicles’ coordinates and quantifies the variations in their respective intensities. Subsequently, the algorithm processes and retrieves pertinent information from large datasets surpassing that of conventional methods under our evaluation, affirming its efficacy. Furthermore, the tool exhibits adaptability for the image analysis of diverse cellular processes, requiring only minimal modifications to ensure accurate detection of exocytosis.

## INTRODUCTION

Intercellular communication facilitated predominantly through the release and uptake of regulatory messenger molecules, is crucial for cell survival. Exocytosis, the process of vesicular release from the plasma membrane, plays a central role in this cellular dialogue. Understanding the regulation of exocytosis at a molecular level has significantly advanced our knowledge of cellular behaviour and its alterations in various disease states (1,2).

Traditional techniques such as electron microscopy and patch-clamp capacitance measurements have provided in-depth insights into exocytosis, albeit without real-time resolution (3–6). The advent of fluorescence microscopy-based imaging, particularly Total Internal Reflection Fluorescence Microscopy (TIRF), along with the advances in 2-D signal processing (7,8), has overcome this limitation, enabling precise visualisation of exocytotic events in a confined spatial dimension and minimising background interference (9,10).

The transition from image acquisition to data interpretation, however, presents challenges due to the voluminous nature of the generated data and the potential for human error in manual analysis. This necessitates an automated approach in the image analysis pipeline. Existing platforms like Fiji, Metamorph and many more, while useful, lack the versatility required for diverse biological analyses. Recent developments have seen the emergence of tools specifically designed for exocytosis event analysis (8,11,12).

Our work diverges from these complex methodologies, aiming instead for a simplified, scalable, and rapid detection of exocytosis. We employ adaptive spatiotemporal filters to identify exocytotic events based on fluorophore activity, followed by trajectory tracking across frames. This approach offers scalability and efficiency, serving as a robust data source for advanced model training, and presents a viable alternative to existing automated analyses of exocytotic events.

## RESULTS

### 1) Image processing algorithm

To process the images for analysis, a scalable and parallel image processing code was developed in MATLAB, which identified the position as well as the gray-scale intensities of the vesicles tagged by fluorophores, captured using TIRF microscopy. The data was subsequently integrated in a Lagrangian particle tracking based macro developed by Crocker and Grier (13) and Boltyanskiy et al. (14) to quantify the intensities of the identified particles and construct particle trajectories. The detailed steps include:

- **Image Pre-Processing:** The raw images (Fig. 1A-sample raw image) were subjected to a series of adaptive spatial filters, enhancing the clarity and focus for subsequent analysis. The intricate details of these processes are elaborated in the methods section (Fig 1B).
- **Binarisation:** We adapted and modified the principles of Otsu’s (15) and Bradley’s algorithms (16), tailoring them to the unique intensity distribution characteristic of TIRF images, thereby ensuring precise detection. Further elaboration is provided in the methods section (Fig 1C).
- **Mask Pre-Processing:** The generated mask underwent a pre-processing scheme, effectively identifying and eliminating areas devoid of cellular material. This refined mask was then applied to the images, assigning a zero-intensity value to regions lacking any cellular data, which significantly streamlined the data size for subsequent analytical steps. Additional details are available in the methods section (Fig 1D).
- **Noise Removal and Filtering:** Utilizing the regional maxima algorithm (17–19), we identified vesicles and their clusters within the processed images. The positions of these entities were determined with an accuracy up to a single pixel. Given the discrete nature of pixel values and the inherent pixel-level quantization of intensities in TIRF images, this accuracy is subject to peak-locking, a phenomenon where the determined positions are constrained to the pixel grid. To overcome this limitation and enhance positional precision, we integrated sub-pixel estimators (20,21) into our algorithm. Drawing from the established efficacy of sub-pixel methodologies in Particle Image Velocimetry (PIV) (22) and Particle Tracking Velocimetry (PTV) for nuanced velocity measurements (20,23,24), our approach aligns with these advanced techniques. The implementation of a sub-pixel routine (21) in our code further refined the accuracy of position determination. A detailed exposition of these methodologies and their application is provided in the methods section (Fig 1-E, H, and I).

**Fig 1:**
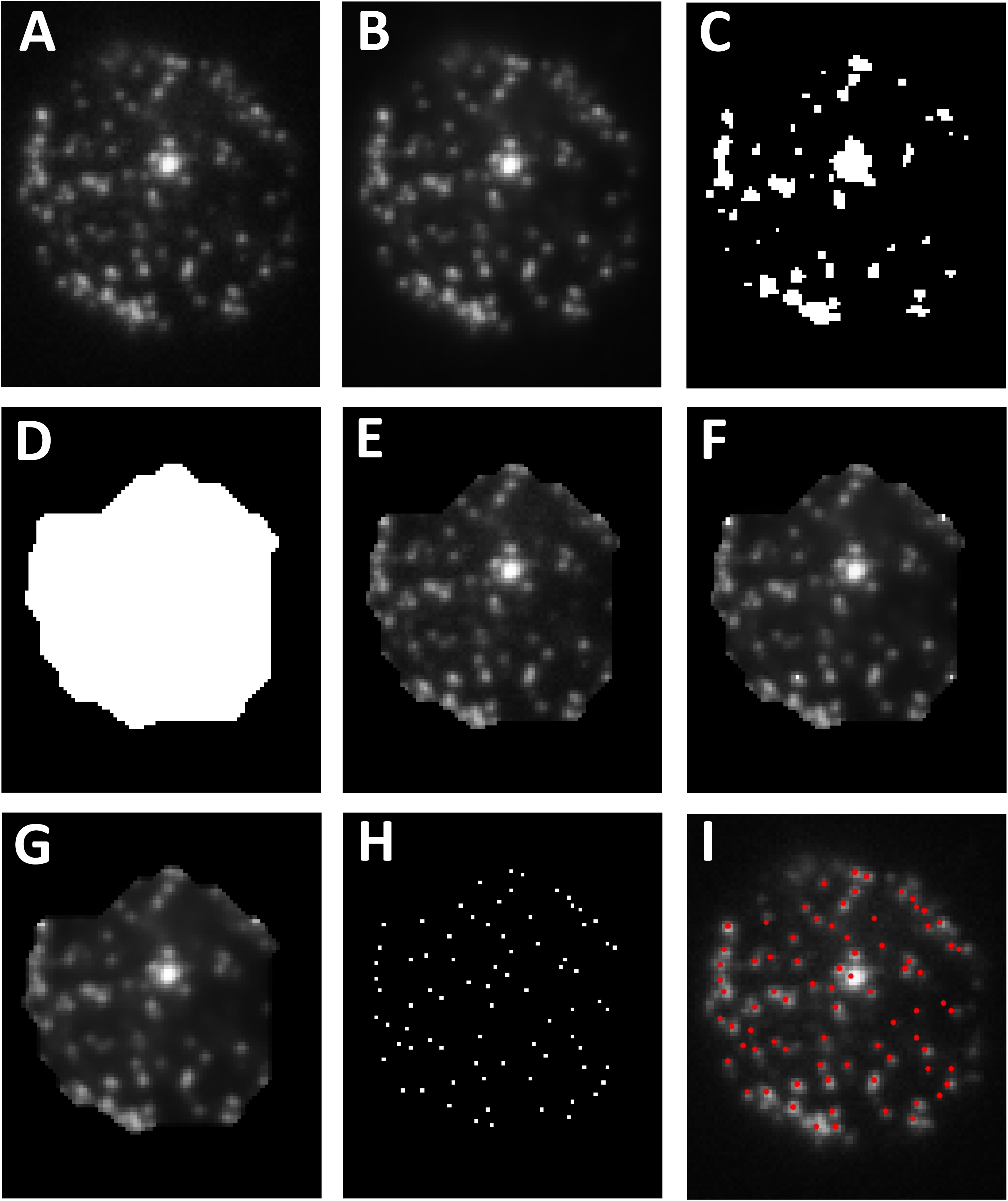
This figure details the various stages in our image processing methodology. A) Displays the original, unprocessed image. B) Shows the image after undergoing preprocessing, which involves the use of an edge preserving filter such as the Weiner filter. C) Illustrates the image post binarization. D) Depicts the image with an improved mask. E) Demonstrates the image with noise eliminated. F) Presents the image after filtering with the help of Weiner, sharpening and Gaussian filters. G) Shows the image post H-Max Transform. H) Displays the image after identifying regional maxima. I) Highlights the particles that have been identified.

These processes ensured that the images captured using TIRF microscopy could be analyzed, the verification of the algorithm is performed at various levels below.

### 2) Validation of the image processing algorithm using artificial images

Primary validation of the algorithm was done with the use of artificial images. Artificial images were generated with 10% signal and 90% noise (Fig 2A and B). This exercise shows the robustness of the code we developed to analyze images with a very low signal to noise ratio. The artificial images were created as described in the methods. A set of five images (n=5) was created, each varying in the level of grey noise introduced. The images were also subjected to analysis using our in-house developed algorithm and compared with two of the tested and accepted methods of particle detection, that is manual counting (described in the methods) and find maxima-based detection (described in the methods) (Fig 2C). The particle counts derived from our algorithm were found to be consistent with those obtained from the manual and find maxima-based methods, yielding a P value of 1 in the One-way Anova comparisons between “by eye” versus “algorithm” and “find maxima” versus “algorithm” (Fig 2D). Notably, the algorithm demonstrated remarkable efficiency in terms of analysis time, significantly (P value – 2.56*10^-39^) reducing the time required for accurate particle detection, as illustrated in Fig 2E.

**Fig 2:**
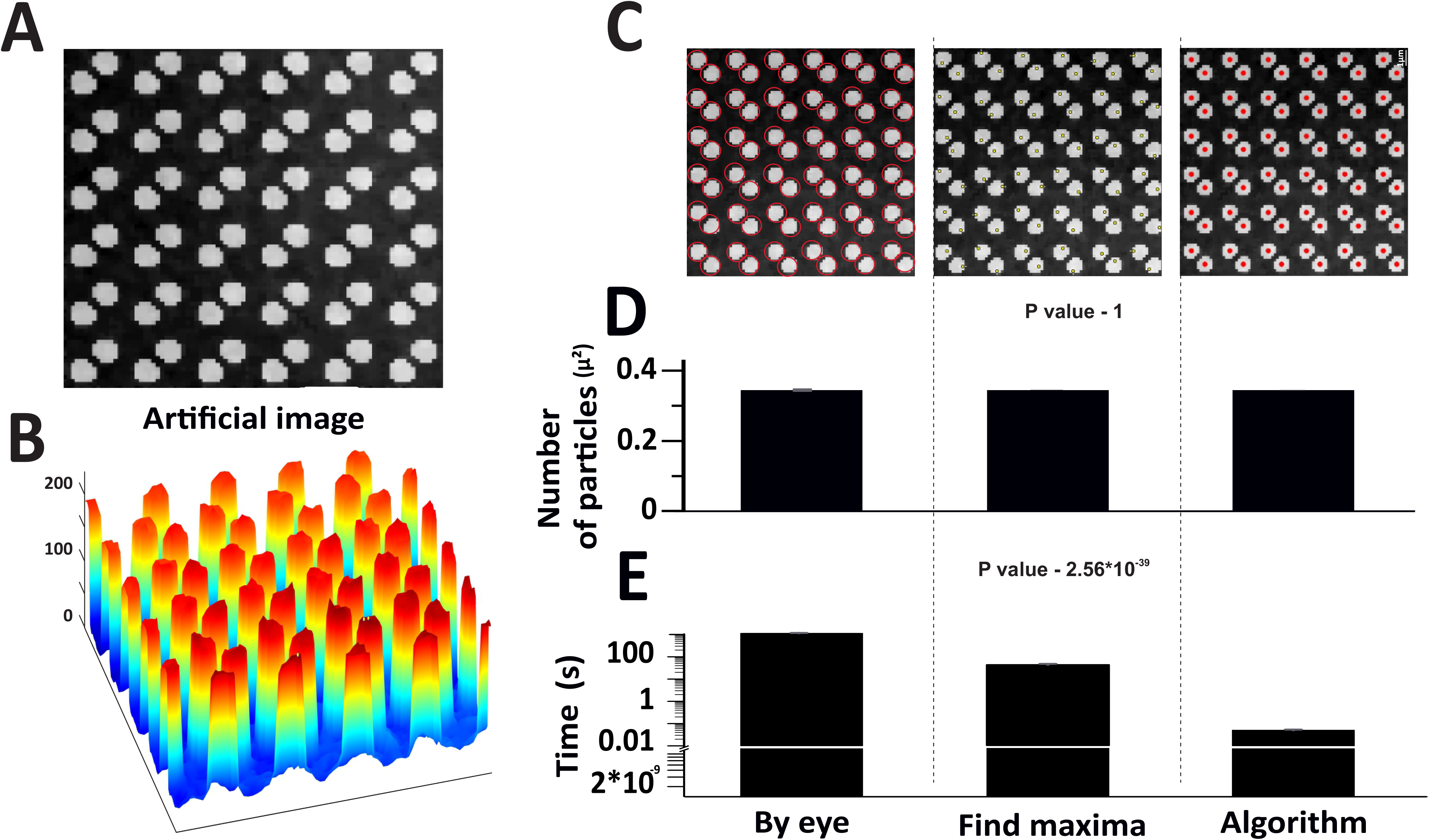
Validation of the image processing algorithm was performed using noisy artificial images that were created with 90% noise and 10% signal. A) Representative artificial image showing the particles. B) Spatial intensity maps of the representative artificial images where the peaks indicate the intensities of the particles C) Detection of particles in noisy artificial images shown visually for all three methods – manual by eye analysis, find maxima function on ImageJ and the image processing algorithm on Matlab. D) Accuracy plot - Average counts per square micron plotted for all three methods. The analysis was performed by three users and averaged, and SEM was calculated. E) Time plot - Average time taken for the analysis was plotted for all three methods. The time taken for analysis was calculated by three users and averaged, and SEM was calculated.

### 3) Verification of the algorithm using beads images

We next challenged our algorithm to analyze images of Tetra spec beads (Fig 3A and B). These are generally used to calibrate fluorescence microscopes, especially for multi color applications. These 0.1 micrometer polystyrene beads, stained with four fluorescent dyes, were prepared and imaged using a 488-nanometer laser as outlined in the methods section. Bead images (n=5) were subjected to analysis by the algorithm (Fig 3C) and compared with 1) manual counting and 2) find maxima-based detection (Fig 3C). The results as shown in Fig 3D, indicate that the algorithm has successfully been able to detect most of the particles (P value – 0.1258). The statistical insignificance also implies the same. Notably, the algorithm showcased exceptional efficiency in terms of analysis time. While manual detection required up to 20 minutes and find maxima-based detection took approximately 20 to 30 seconds, our algorithm significantly expedited the process (P value – 2.24*10^-40^), as illustrated in Fig 3E.

**Fig 3:**
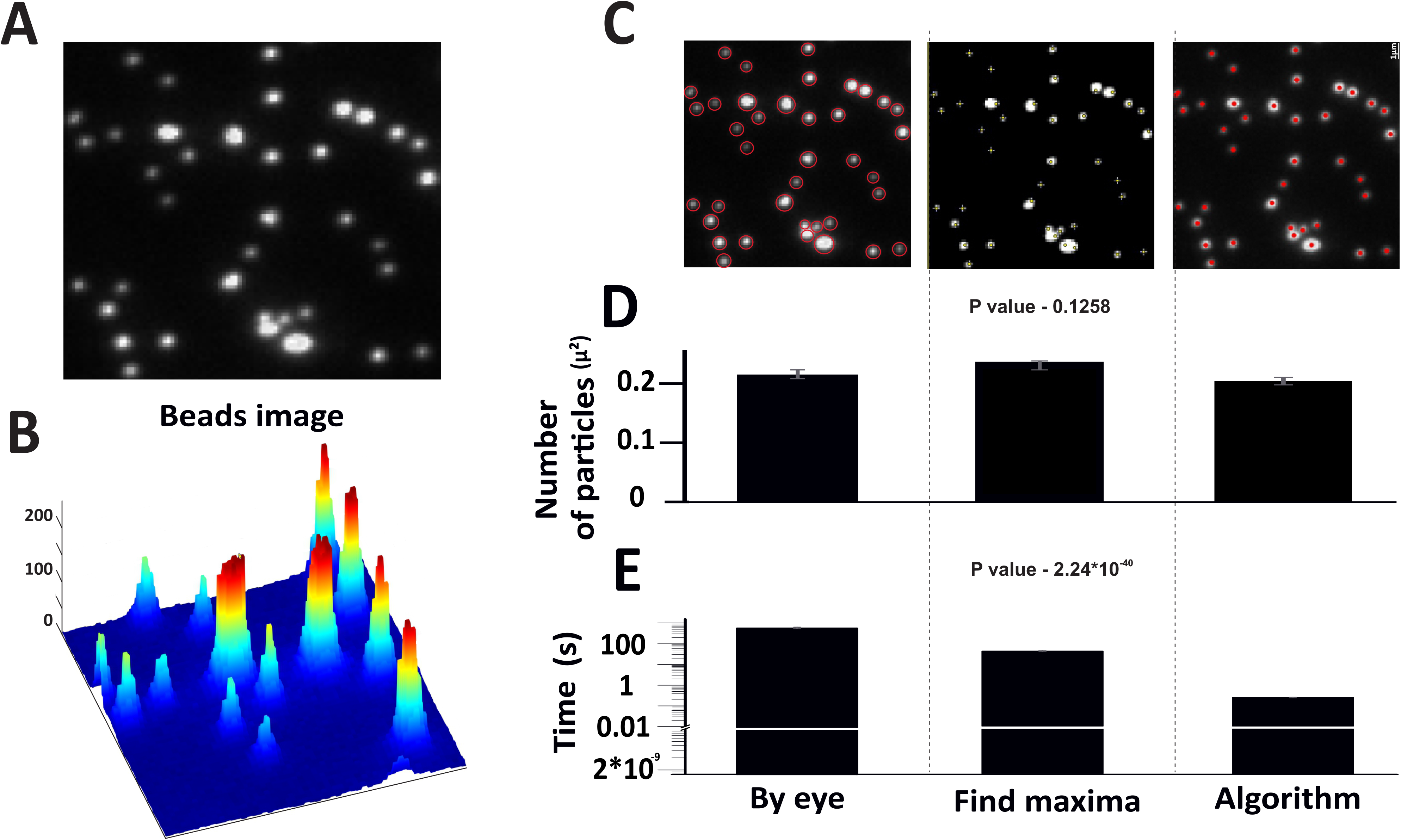
The efficiency of the image processing algorithm was evaluated using images of tetraspeck beads. A) The image represents tetraspeck beads imaged using TIRF microscopy. B) The peaks in the spatial intensity maps of the artificial images signify the intensities of the particles. C) Three methods were employed to analyze beads images: manual eye analysis, the find maxima function on ImageJ, and an image processing algorithm on Matlab. D). The accuracy of the three methods is plotted showing the average counts per square micron in each case. The analysis was performed by three users, and the results were averaged, with the standard error of the mean (SEM) calculated. E). A time plot was also generated, showing the average time taken for the analysis for each method. The time taken for analysis was calculated by three users and averaged, with the SEM.

### 4) Analysis of TIRF images of mammalian cells

Mammalian cells in culture are transfected or transduced as mentioned in the methods and imaged using TIRF microscopy (Fig 4A and B). These images are more complex with respect to the low signal to noise ratio that arises due to the inherent fact that different cells express the fluorescent molecules at different levels and hence the disparity. The signal seen in these images are from the large dense core vesicles tagged to a fluorophore (details in the methods). Images (n=5) were subjected to analysis using the algorithm and it was able to almost detect many numbers of particles which was compared with the manual and find maxima-based detection (P value – 0.9941) (Fig 4C). The accuracy of detection is quite similar as plotted in Fig 4D and shows the same with statistical insignificance. Like the previous trend with artificial images and beads, the algorithm took milliseconds when compared to seconds using find maxima and minutes by manual detection (P value – 4.2*10^-12^) (Fig 4E).

**Fig 4:**
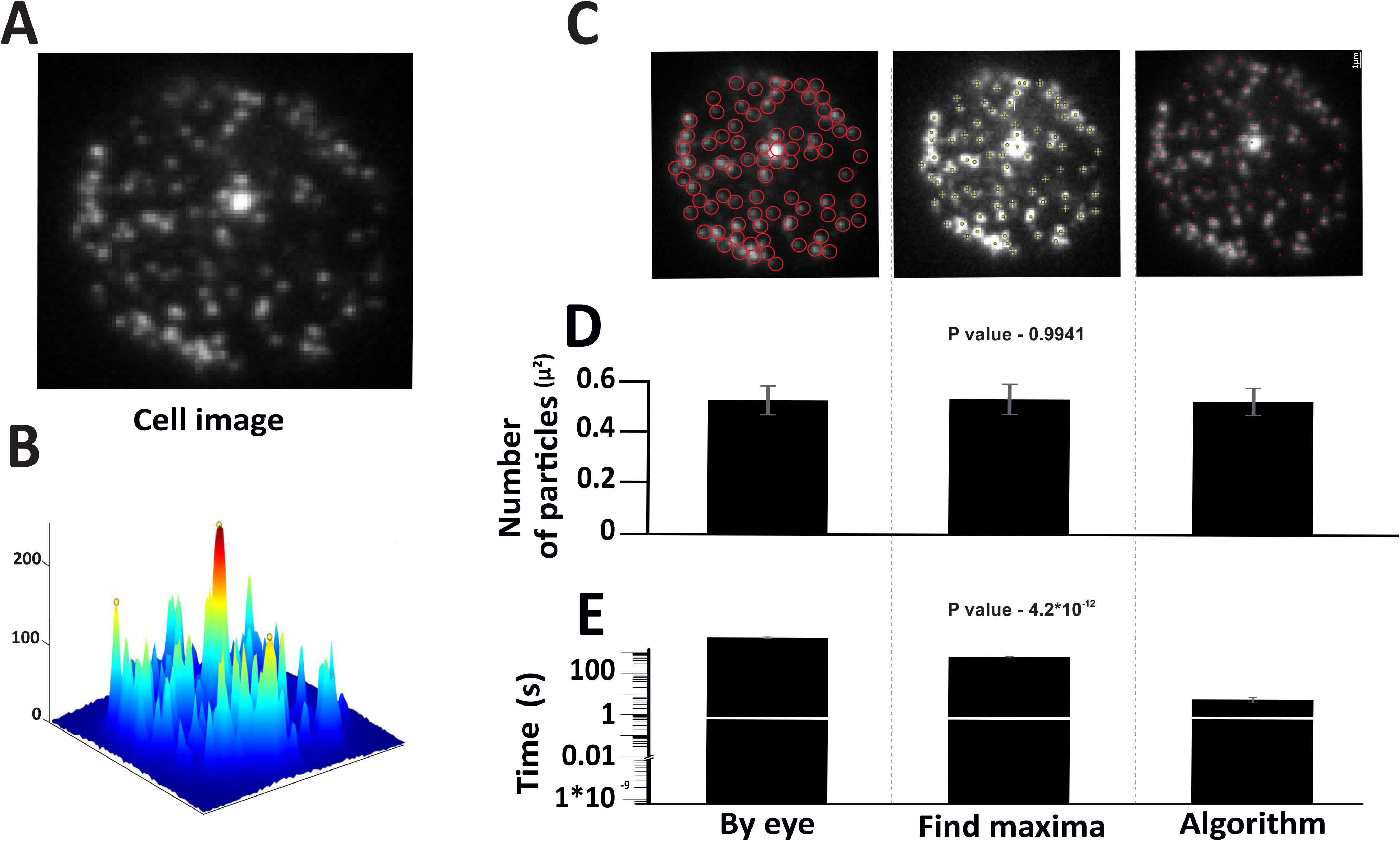
The ability of the image processing algorithm to identify Large-dense core vesicles was put to test. A) Indicates fluorescently labelled cell images. B) Spatial intensity maps showing the vesicles identified. C). Images were subjected to analysis by three methods – manual by eye analysis, find maxima function on ImageJ and the image processing algorithm on Matlab. D) Accuracy plot - Average counts per square micron were plotted for all three methods. The analysis was performed by three users and averaged, and SEM was calculated. E) Time plot - Average time taken for the analysis was plotted for all three methods. The time taken for analysis was calculated by three users and averaged, and SEM was calculated.

### 5) Detection of exocytosis by the algorithm

Next the idea was to analyze images of dynamic cellular processes using the algorithm. A detailed description of how the algorithm performs this task is explained as a part of the first result. Fast events such as regulated exocytosis have always been a challenge to analyze since it is time-consuming. Exocytosis is marked by the sudden peak in the intensity followed by loss of intensity. In our previous work, we analyzed such events manually using Metamorph (to move between different frames) where the exocytosis events were marked. Such events were now subjected to the analysis using the algorithm. The first step in both the modes of analysis is to mark or detect the vesicles (Fig 5A). For a closer view, a montage of one of the vesicles’ undergoing exocytosis is shown in Fig 5B. This particle has been marked using a white square in Fig 5A. The loss of intensity after the first few frames of the montage is clearly visible. This loss of intensity indicates the phenomenon of exocytosis. Spatial intensity maps shown in Fig 5C also reinforce the loss of intensity in frame 586 whereas this particle was seen in frame 580. Further, intensity versus time plots were generated for this particular particle in Metamorph (Fig 5D) and using the algorithm (Fig 5E). These plots clearly indicate the sudden drop in the intensity at frame 586 indicating exocytosis. Another highlight is the fact that the algorithm is also especially robust in determining the exact frame number or time point at which the exocytosis of the vesicle is seen.

**Fig 5:**
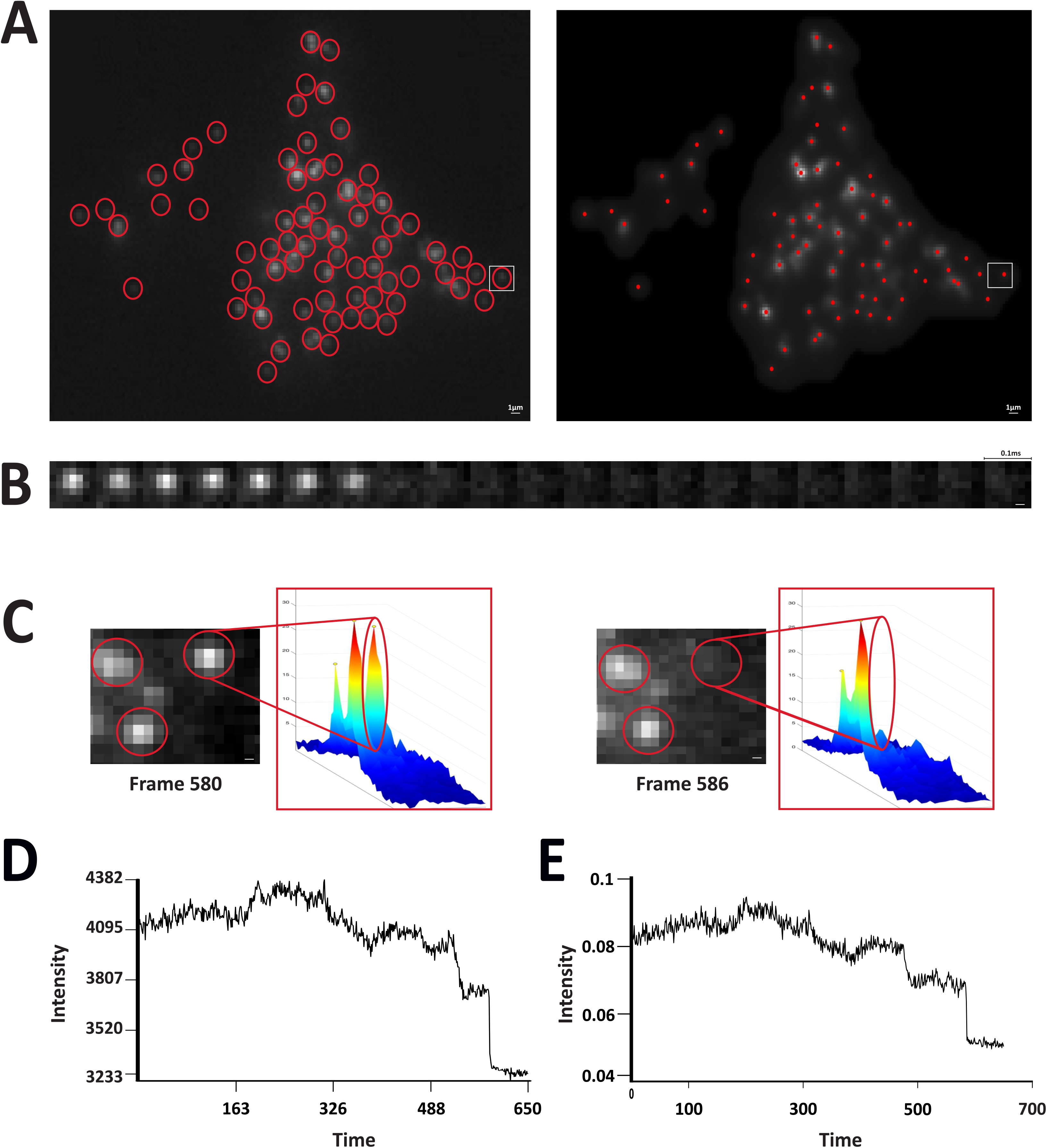
Testing the ability of the algorithm to detect exocytosis events. A). Snapshot of the movie, where the particles are identified manually using Metamorph and by the algorithm. The particle highlighted with a white square is the one undergoing exocytosis. B). An image sequence generated using Metamorph showing the K^+^ stimulated exocytosis of the particle marked with a white square in Fig A. C) A small part of the whole cell image at frame 580 and 586 and the corresponding spatial intensity maps. It indicates the loss of the particle at frame 586 with the absence of an intensity peak marked with an empty red oval. D & E). Intensity vs time plots obtained using Metamorph and the Algorithm. The intensity of the particle over the entire movie is depicted by these plots. A sudden dip in intensity is seen at frame 586-590 as shown by the plots generated by both manual analysis and the algorithm.

## METHODS

### Cell culture

#### Human Pancreatic Islet Cells

Pancreatic islets were obtained from human cadaveric donors by the Nordic Network for Clinical Islet Transplantation (ethical approval by Uppsala Regional Ethics Board 2006/348), the ADI Isletcore at the University of Alberta (ethical approval by Alberta Human Research Ethics Board, Pro00001754 or the Institutional Human Ethics Committee, Indian Institute of Science, 08/20.07.2022), with written donor and family consent for use in research (25). Work with human tissue complied with all relevant ethical regulations for use in research and the study was approved by the Gothenburg Regional Ethics Board, Sweden and ethical committees at Indian Institute of Science, India.

CMRL 1066 culture medium containing 5.5mM glucose, 10% fetal calf serum, 2mM L-glutamine, streptomycin (100U/ml), and penicillin (100U/ml) was used to culture the isolated islets at 37°C in an atmosphere of 5% CO_2_ for up to two weeks. Prior to imaging, islets were dispersed into single cells using Ca^2+^-free cell dissociation buffer (Thermo Fisher Scientific) supplemented with containing 10% (v/v) trypsin (0.05% Thermo Fisher Scientific) and gentle agitation. Dispersed cells were sedimented by centrifugation and resuspended in serum-containing medium before plating onto 22-mm Poly-L-Lysine-coated coverslips and allowed to settle overnight. Seeded cells were infected using adenovirus coding for granule markers (NPY-Venus or NPY-mCherry) and imaged 24-36 hr later. In some cases, cells were in addition transfected with cDNA coding for EGFP-labelled proteins at the time of infection. Transfection was done using Lipofectamine 2000 (Thermo Fisher Scientific) according to the manufacturer’s protocol.

#### Beads

100nm tetraspeck beads (Thermo Fisher Scientific) were diluted in a 1:100 ratio using PBS. They were immobilized on the coverslip surface/ a glass bottom dish and imaged.

#### Artificial images

To rigorously test and validate our image processing algorithm, we engaged in the generation of artificial images. The creation of circular objects within these images was guided by the locus of a circle, defined mathematically by the equations:

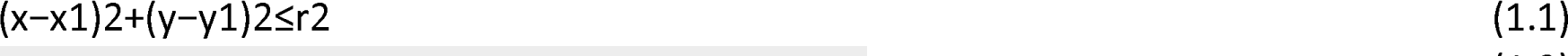

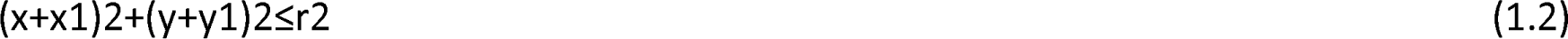

Here, (x1, y1) represents the circle’s centre coordinates, and (r) denotes its radius. This formulation yields two distinct circular regions. By applying a logical ’or’ operator, we derived a region belonging to either of the two circular areas defined by equations (1.1) or (1.2).

The radius (r) was adjustable, allowing for the simulation of various vesicle regions encountered in TIRF images, ranging from intersecting at two points, one point, or not intersecting at all. This pattern was then replicated across the image to generate multiple circular regions.

To introduce realistic noise into the image, we utilized multiple noisy images. The pixel values of both the raw and noisy images were scaled between 0 and 1. Their linear combination, as depicted in equation (1.3), resulted in the final noisy image:

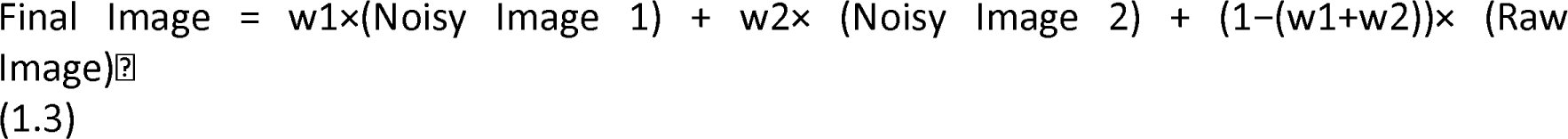

Final Image = w1×(Noisy Image 1) + w2× (Noisy Image 2) + (1−(w1+w2))× (Raw Image)  (1.3)

#### Buffers and solutions

Cells were imaged in (mM) 138 NaCl, 5.6 KCl, 1.2 MgCl_2_, 2.6 CaCl_2_, 3 or 10 D-glucose, 5 HEPES (pH 7.4 with NaOH) at ∼32 °C. Exocytosis was evoked with high 75 mM K^+^ (equimolarly replacing Na^+^) or 10 mM glucose, applied by computer-timed local pressure ejection through a pulled glass capillary. For K^+^-induced exocytosis, spontaneous depolarizations prevented with 200 μM diazoxide; this protocol depolarizes the cells to zero mV within ∼50 ms. forskolin (Fsk; 2 μM) was included.

#### Microscopy

Cells and beads were imaged using a total internal reflection (TIRF)/ Confocal microscope based on an AxioObserver Z1/Nikon Ti2 Eclipse with a 100x/1.45 objective (Carl Zeiss/Nikon). Excitation was done by two DPSS lasers at 491/488nm. The emission light was captured on a EMCCD camera (Photometrics/Andor). Scaling was maintained at 160/130 nm per pixel.

### Analysis

#### Processing of images

The first frame of the cells and beads movies was duplicated and saved after applying the median filter in Fiji. Artificial images were also analyzed after the application of the median filter.

#### By eye analysis

The counting of the particles was performed by three independent users on Metamorph. The output was averaged, and SEM was calculated. The counting was timed, and the average time was calculated.

#### Find maxima

Granule density was calculated using a script that used the built-in ‘find maxima’ function in ImageJ (http://rsbweb.nih.gov/ij) for spot detection. Counting was done by 3 independent users. Three independent users performed the particle counting, and their results were averaged to calculate the SEM. Additionally, the time it took to complete the counting was recorded and averaged.

#### Algorithm

3 independent trails were performed using the algorithm. The output was averaged, and SEM was calculated. The analysis was timed, and the average time was calculated and averaged, followed by the calculation of SEM.

#### Image processing algorithm

The following are the detailed steps involved in the working of this algorithm.

##### i. Image Pre-processing

At the outset, the images undergo a pre-processing phase. This phase employs the use of a Weiner filter. This particular filter is adaptive and edge-preserving, enhancing the signal-to-noise ratio (SNR) in the image, which is crucial for the clarity and accuracy of subsequent processing steps (refer to Fig 1B for a visual representation).

##### ii. Binarization

Once pre-processing is complete, the images are binarized. Binarization is the process of converting an image into a binary format, which essentially means it will have only two possible values for each pixel. The challenge here is to determine the threshold for this conversion. To automate this process and enhance efficiency, the renowned Otsu’s Algorithm from 1979 is employed. This algorithm is particularly adept at evaluating the threshold by utilizing the zeroth and first-order cumulative moments. However, a limitation of Otsu’s Algorithm is its inability to account for spatial variations in intensity. To address this, the Bradley’s Adaptive Thresholding Algorithm from 2007 is also considered. While this algorithm is sensitive and accounts for spatial variations, it can sometimes be overly sensitive for certain images, as observed in Supplementary fig 1.

To strike a balance and ensure optimal thresholding, the image is divided into smaller sub-images, each measuring 32 X 32 pixels. Otsu’s Algorithm is then applied to each of these sub-images. The resultant image, which can be seen in Fig 1C, was further refined. Any non-detection of low-intensity bright pixels was addressed by either adopting a smaller sub-image size of 16 X 16 pixels or by adjusting the value obtained from Otsu’s Algorithm for specific sub-images.

#### iii. Pre-Processing of Mask

The next phase involves refining the mask generated in previous step, so that it encompasses a specific cluster of cells. This is crucial for isolating areas of interest within the image. The mask generation employs morphological closing operations, which involve dilation followed by erosion using the same structuring element. The dilation process adds pixels to object boundaries, while erosion removes them. The choice of structuring element is pivotal, and given the shape of the cells, a disc structuring element was deemed most appropriate. To bridge larger gaps, the disc’s radius was set at a relatively higher value of 100 pixels. Subsequent operations were performed to refine the mask, including dilation with a disk of radius 5 ± 3 pixels and an opening operation with a disk of radius 3 ± 1 pixels. The sequence of these operations transformed the image from Fig 1C to Fig 1D.

#### iv. Noise Removal and Filtering

With the mask ready, it’s overlaid onto the original raw image (as seen in Fig 1A). This step isolates the cell clusters and assigns zero values to all background pixels, resulting in the image depicted in Fig 1E. The intensity values of this image are then scaled between 0 and 1. The image matrix is visualized as a surface, with intensity values representing the height at each (x,y) position. This topological representation, detailed in Fig 2, allows for the identification of vesicles as local peaks. The Regional Maxima finding algorithm is employed to pinpoint these peaks. However, due to the sensitivity of this algorithm, especially in noisy images, a series of filters, including the Wiener filter, sharpening filter, and Gaussian filter, are applied in succession, before applying regional maxima. If over-segmentation persists, the H-Max Transform is used to suppress smaller peaks, followed by the application of the regional-maxima algorithm. The final positions of these maxima correspond to the vesicle positions, as illustrated in Fig 1I.

#### v. Detection of Dynamic Events

The final stage is centered on detecting dynamic cellular events. This requires identifying particle positions across all images in a stack. These positions are then used to calculate intensity values, taking the average intensity at the particle position and its four neighbouring points. This data feeds into the Tracker, which constructs particle trajectories over time, revealing the position and intensity of vesicles as they move within the cell. In instances where there’s no significant motion, the tracker ceases operation. For such cases, the images in the sequence without significant motion are identified, and their first frame is considered. Intensity values for subsequent frames are calculated based on the vesicle positions determined in the first frame. These intensities are then plotted over time, and to enhance visualization, surface plots are created, with the height of the surface corresponding to the intensity values.

### Statistics

Data are presented as mean±s.e.m, unless otherwise stated. Statistical significance was assessed using One way-Anova, as appropriate. The P values have been included in the figures and the results.

## DISCUSSION

The algorithm we have developed stands out as a rapid and straightforward solution for exocytosis detection, demonstrating high fidelity in particle identification across various image types. Initially, the algorithm’s performance was validated using artificial images, where it successfully identified particles with a high degree of accuracy, aligning closely with the predetermined counts used to create these images. Moving a step further towards real biological samples, the algorithm showcased its proficiency in detecting tetraspeck beads, yielding results comparable to manual and find maxima-based analyses. In the subsequent phase of evaluation, the algorithm was applied to primary tissue samples, specifically β cells of the islet, where it effectively detected fluorescently labelled vesicles, producing counts consistent with those obtained through other analysis methods.

A standout feature of this algorithm is its rapid processing speed, completing analyses in just a few milliseconds and providing valuable data promptly. This efficiency is crucial when dealing with extensive image datasets and time-lapse movies, ensuring that the algorithm meets the demands of high-throughput image analysis.

Pre-processing stands as a pivotal component in our image analysis workflow, addressing the unique challenges posed by each image due to variations in sample preparation, imaging parameters, and other factors. As depicted in Fig 1, our algorithm employs a foundational schematic that adapts to these variations, particularly in the pre-processing stages. This adaptability is achieved through the use of adaptive spatial filters and varying kernel dimensions, tailored to meet the specific requirements.

This approach enhances the algorithm’s capability to handle the complexities found in TIRF (Total Internal Reflection Fluorescence) images, as detailed in the methods section. Despite the variations in spatial filters and their sequencing, the algorithm maintains a consistent procedural structure across different scenarios, as illustrated in Fig 1. By finely tuning these parameters, the algorithm’s accuracy can be elevated, surpassing the 99% threshold validated in our study.

Comparatively, some existing algorithms in this domain utilize Fourier transform-based particle detection strategies, requiring minor modifications to accommodate different image qualities. Our algorithm’s strength lies in its versatility, performing well across a diverse range of images, particle sizes, and noise levels. Many algorithms rely on temporal tracking of identified particles for exocytosis detection. Our algorithm introduces a novel approach, where it tracks the movement of particles based solely on intensity changes and spatial coordinates, potentially reducing the likelihood of false positive detections.

The applications of our algorithm extend beyond exocytosis detection, providing a comprehensive framework for particle identification. This study highlights its utility in studying biological processes, specifically exocytosis, but its potential applications are vast, ranging from co-localization studies to particle tracking and beyond.

**Supplementary fig 1:**
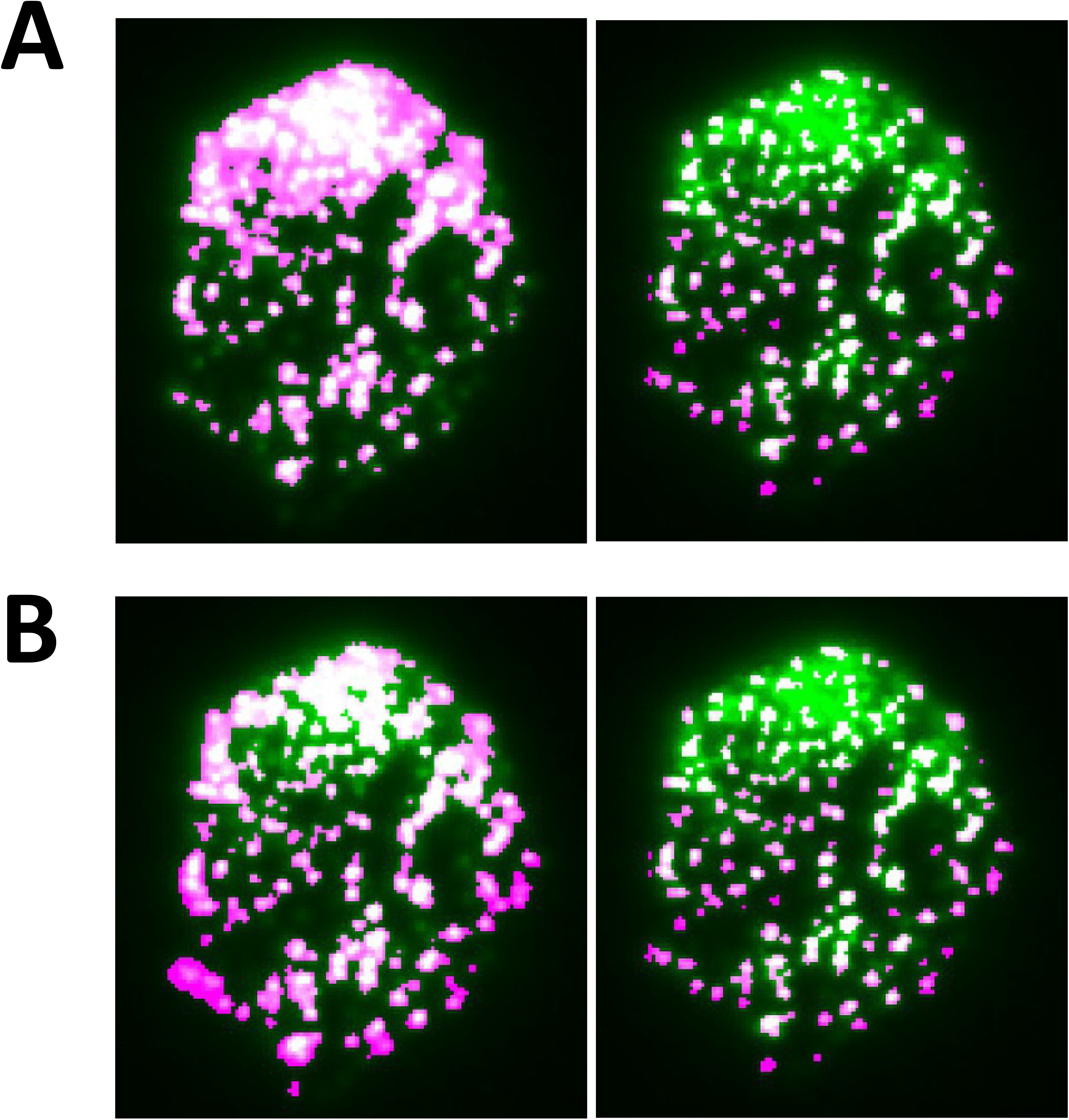
Image depicting the differences in the detection using the developed algorithm in comparison with Otsu’s and Bradley’s algorithms. A) A cell image with punctate structures was used for detection using both the Otsu’s algorithm and our newly developed algorithm. This depicts the inability of Otsu’s algorithm to take into consideration the spatial variations in intensity. B) Bradley’s algorithm was used to account for spatial variations, but it turns out to be overly sensitive.

**Supplementary fig 2:**
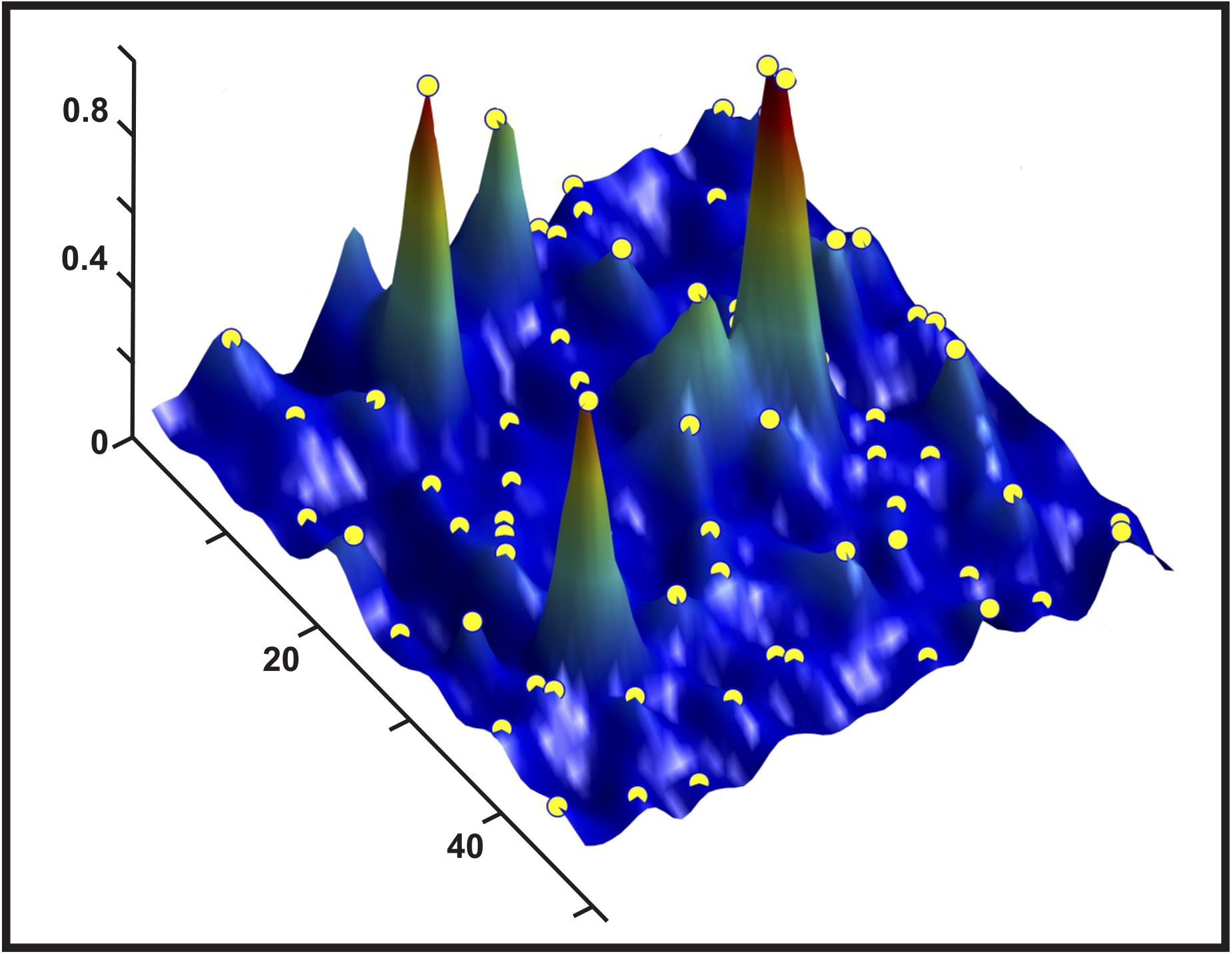
Surface Plot of an Image. The yellow spots correspond to the regions which could potentially contain punctate structures or vesicles.

## Author Contributions

Conceptualization – AAM, AD, SM and NRG; Methodology - AAM, AD, SM and NRG; Software – AD and SM ; Validation - AAM, AD, SM and NRG, ; Formal analysis – AAM and AD,; Investigation AAM and NRG,; Resources - NRG,; Data curation - AAM, AD, SM and NRG; Writing - AAM, AD and NRG; Review & editing - AAM, AD, SM and NRG, ; Supervision – SM and NRG,; Project administration – AAM and NRG, ; Funding acquisition - NRG. All authors have read and agreed to the published version of the manuscript.

## Acknowledgments

We would like to thank Vibha V, Sayoni Maiti, Anuma Pallavi and Veerabhadraswamy Priyadarshini, for helping us during the analysis and for giving their valuable insights.

